# Environmental factors shape the culturable bacterial diversity in deep-sea water and surface sediments from the Mariana Trench

**DOI:** 10.1101/2022.01.13.476188

**Authors:** Yuewei Ma, Wenmian Ding, Yuepeng Wang, Ping Chen, Hui Zhou, Xuan Li, Yanyan Huang, Peng Nan

**Affiliations:** Ministry of Education Key Laboratory for Biodiversity Science and Ecological Engineering, School of Life Sciences, Fudan University, Shanghai, China; CAS-Key Laboratory of Synthetic Biology, CAS Center for Excellence in Molecular Plant Sciences, Institute of Plant Physiology and Ecology, Chinese Academy of Sciences, Shanghai, China; University of Chinese Academy of Sciences, Beijing, China; Chinese Ancient Books reservation and Conservation Institute, Fudan University, Shanghai, China; Fudan Zhangjiang Institute, Shanghai, China

**Author notes:** Contributed equally (Yuewei Ma, Wenmian Ding).

**Keywords:** Mariana Trench, deep-sea sediment, culturable microbial diversity, environmental parameters, Mantel test

## Abstract

Hailed as “The Fourth Pole”, the Mariana Trench is the deepest part of the ocean. The microbial diversity in it is extremely complicated, which might be caused by the unique environmental factors such as high salinity, low temperature, high hydrostatic pressure, and limited nutrition. Based on 4 seawater samples and 4 sediment samples obtained from the Mariana Trench, we isolated and fostered the microorganism clones with kinds of culture mediums and high-throughput culturing. By using the molecular identification methods based on PCR of 16S rDNA and ITS gene, 1266 bacterial strains in total were isolated and identified, which affiliated to 7 classes, 16 orders, 25 families and 36 genera in four phyla : Proteobacteria, Firmicutes, Actinobacteria and Bacteroidetes. Strains in genera *Halomonas, Pseudoaltermonas* were the dominant bacteria isolated from the samples. With Mantel tests on the sample-environmental parameter matrix, the sample-environmental organic matter diversity matrix and the sample-microbial diversity matrix, we concluded that the environmental parameters and the organic matters in the condition can shape the culturable bacterial diversity in deep-sea water and surface sediments from the Mariana Trench.

## Introduction

The deep sea, generally defined as the ocean below 1000m, accounts for 75% of the total volume of the earth’s oceans. The average depth of the ocean is 3800m, and the deepest trench is about 11000m. The deep-sea environment has the following general characteristics: darkness, low temperature, high pressure, high salinity and oligonutrition. The oligotrophic condition makes the deep-sea microorganism have a high potential for metabolic research.

The vertical distribution of nutrients in the Mariana Trench shows that the temperature of the deep-sea water decreases slowly with the increase of the depth, while the salinity rises slowly with the increase of the depth until it stabilizes at about 34.7, and the dissolved oxygen, nitrogen and sulfur contents also show a decreasing trend below 1000m of the sea level. The nutrient concentration in the surface of the sediment is slightly higher than that in the deep sea, but it decreases with the increase of the depth in the sediment. The dissolved oxygen and nutrients in seawater and sediment of Mariana Trench are all at a low level, while the deep-sea habitats show obvious vertical distribution. The unique species diversity and genetic diversity of deep-sea microorganisms are caused by the oligotrophic environment. In order to adapt to the extreme environment, the microorganisms living in the deep-sea may produce bioactive metabolites with great application potential. There are also studies showing that deep-sea microbes can use refractory organic matter more efficiently than those at the surface.

With the development of 16S rRNA sequencing, SSU rRNA sequencing and metagenomic sequencing technologies, researchers have gained some understanding of the community structure of microorganisms in the Mariana Trench, but there is still a large gap in the isolation and culture of microorganisms in the Mariana Trench. The isolation of culturable microorganisms is essential for the study of their metabolic potential, functions, applications. Little research has been done on culturable microbes, and isolation specific to the microbes in the deep sea environment is still in its infancy.

To obtain pure cultured strains is the prerequisite for studying the adaptive evolution, metabolic regulation and application and development potential of deep-sea microorganism. Therefore, it is necessary and urgent to study the isolation, culture and identification of microorganisms in the deep-sea sediments of Mariana Trench.

## Materials and Methods

### Sampling and cultivation

Seawater and sediment samples were collected from the Challenger Deep of the Mariana Trench. Information of the collection sites were shown in Table 1. The seawater samples were diluted at three gradients of 10^−1^, 10^−2^ and 10^−3^, while the surface sediment samples were diluted at seven gradients from 10^−1^ to 10^−7^. 200μL of each dilution was plated on 6 different media (Table S1) separately and incubated at 28 °C for 3-7 days. Serial dilutions and plate counting were used to calculate the microbial abundance in the 8 samples.

**Table 1.** Geochemistry parameters of the studied sites in the Mariana Trench.

| Sample | Sampling position | Sampling date | Longitude | Latitude | Depth | Sample Type |
| --- | --- | --- | --- | --- | --- | --- |
| S1 | MBR04 | 2019.12.05 | E142.23° | N10.82° | 5842 | sediment |
| S3 | MBR05-A1 | 2019.12.09 | E141.95° | N10.98° | 6957 | sediment |
| S2 | MBR06-A1 | 2019.12.11 | E141.18° | N10.81° | 6477 | sediment |
| S4 | S4 | 2018.12.12 | E142.20° | N11.33° | 10900 | sediment |
| W1 | MBR04 | 2019.12.07 | E142.23° | N10.82° | 1000 | seawater |
| W2 | MBR04 | 2019.12.07 | E142.23° | N10.82° | 4000 | seawater |
| W3 | MBR02 | 2019.11.24 | E142.19° | N11.33° | 5900 | seawater |
| W4 | S4 | 2018.12.12 | E142.20° | N11.33° | 10900 | seawater |

### Identification of microorganism

After 3-7 days’ plate culture, the cultured single colonies were picked up for individual colony PCR. The single colonies were subjected to PCR approach with the bacterial 16S rRNA sequence general primers 27F/1492R.The PCR system was composed of 12.5μL 2xTaq PCR Mastermix, 0.5μL 8F primer, 0.5μL 1492R primer and 11.5μL ddH2O. The PCR procedure was as follows: pre-denaturation at 94 ° C for 3min, denaturation at 94 ° C for 30s, annealing at 52 ° C for 30s, extension at 72 ° C for 90s, and 30 cycles. Elongation at 72°C for 10min.The PCR products were separated by 1% agarose gel electrophoresis, and the single target bands were further sequenced.

### Sequencing and phylogenetic analysis

The sequences were blasted against NCBI database, and similarity comparison analysis was conducted on the microbial species information. The 16S rRNA gene sequences of 1 to 2 highly homologous model strains of each genus were selected to construct the N-J evolutionary tree using MEGA X. The phylogenetic matrix was estimated according to the Kimura model with Bootstrap value of 1000 times.

### Analysis of environmental organic compounds by LC-MS

50 mg sediment sample were accurately weighed. The metabolites were extracted using a 400 μL methanol : water (4:1, v/v) solution with 0.02 mg/mL L-2-chlorophenylalanin as internal standard. The mixture was allowed to settle at −10°C and treated by High throughput tissue crusher Wonbio-96c at 50 Hz for 6 min, then followed by ultrasound at 40 kHz for 30 min at 5°C. The samples were placed at −20 °C for 30 min to precipitate proteins. After centrifugation at 13000g at 4 °C for 15 min, the supernatant were carefully transferred to sample vials for LC-MS/MS analysis.

The mobile phase was composed of 0.1% formic acid in water : acetonitrile (95:5, v/v) (solvent A) and 0.1% formic acid in acetonitrile : isopropanol : water (47.5:47.5:5, v/v) (solvent B). The sample injection volume was 2 μL and the flow rate was set to 0.4 mL/min. The column temperature was maintained at 40 °C.The tandem mass spectrometer was operated in positive mode electron spray ionization (ESI) and the parameters were shown in Table S2.

### Statistical analysis

The R package VEGAN (v.4.1.0; R Foundation for Statistical Computing, Vienna, Austria, https://www.r-project.org) was used to construct the distance matrix for the *sample-metabolite diversity matrix* with Bray-Curtis distance, and the distance matrix for the *sample-microbial diversity (at order level) matrix* with euclidean distance, then the mantel test is performed on the two distance matrices.

The R package UpSetR (v.4.1.0; R Foundation for Statistical Computing, Vienna, Austria, https://www.r-project.org) was used to draw the upset plot. The R package pheatmaps (v.4.1.0; R Foundation for Statistical Computing, Vienna, Austria, https://www.r-project.org) was used to draw the correlation heatmaps.

## Acknowledgement

All 16S rDNA sequences in this study were submitted to the NCBI database. The data of LC-MS untargeted metabolomics were analyzed on the free online platform of Majorbio Cloud Platform (https://www.majorbio.com).

## Results

### Quantification of microbes at different stations

Microbial abundance was calculated by plate counting method. The abundance of microbes cultivated in the 4 seawater samples was generally of 10^4^, whereas the one in the 4 sediment samples was in the range of 10^6^ to10^7^. Seawater samples W1, W2, W3, which collected from 1km, 4km, and 5.9km below the sea level, showed an increase in colony abundance with increasing depth. The increasing trend and the microbial abundance was consistent with the previous study. The abundance of microorganism isolated from CDB+ was the highest, while the one isolated from MG-was the lowest (Figure 1A).

**Figure 1.**
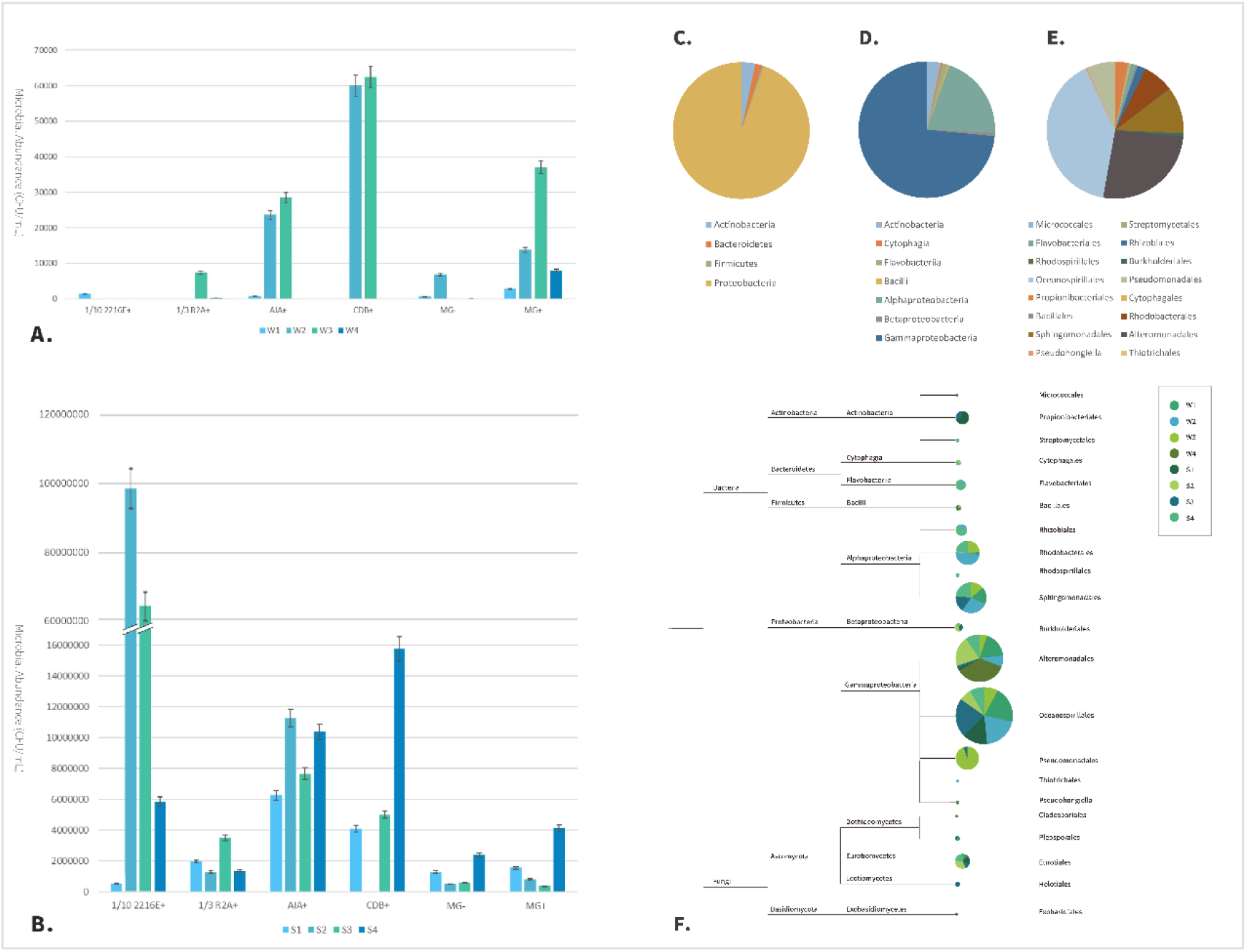
Microbial abundance and diversity of bacteria isolated from the Mariana Trench. (A) Microbial abundance (CFU/ mL) isolated from seawater in different media at 4 different stations in the Mariana Trench; (B) Microbial abundance (CFU/ mL) isolated from surface sediments in different media at 4 different stations in the Mariana Trench; (C-E) Community composition of culturable bacteria from the Mariana Trench (C. at phylum level; D. at class level; E. at order level); (F) The species level tree of the microorganisms isolated from the samples. (The area size of the pie chart shows the number of microorganisms isolated from the order, and the different colors represent the proportion of microorganisms from each site.)

For the 4 sediment samples, there weren’t a general trend of increasing colony abundance with increasing depth. The highest abundance of microorganisms was in 1/10 2216E+ medium, which was far more than other media. The abundance of microorganism isolated from MG+ and MG-was the lowest (Figure 1B).

### Diversity of bacteria isolated from the Mariana Trench

With high-throughout culturing, among the 1266 strains of bacteria with reliable 16S rDNA sequence, a total of 4 phyla, 7 classes, 16 orders, 25 families and 36 genera were obtained.

The highest proportion of the phylum level was Proteobacteria, accounting for 95%, followed by Actinobacteria, 3%, Bacteroidetes, 2%, and Firmicutes accounting for less than 1% (Figure 1C). At the class level, the largest proportion was γ-Proteobacteria (74%), followed by α-Proteobacteria (20%), followed by Actinobacteria (3%), Flavobacteria, β-Proteobacteria, Bacillus and Cytophagia, each accounting for about 1% (Figure 1D). At the order level, the top five orders were Oceanospirillales, Alteromonadales, Sphingomonadales, Rhodobacterales and Pseudomonadales.

#### Diversity of culturable bacteria from different samples

In each sample, γ-Proteobacteria accounted for the largest proportion, ranging from 45% to 97%. α-Proteobacteria were distributed in 6 samples, and accounted for the second largest proportion in 5 samples. In the 4 seawater samples, the main culturable bacteria were γ-Proteobacteria and α-Proteobacteria. While in the four sediment samples, Actinomycetes and Flavobacteriia showed a relatively higher percentage than in seawater samples (Figure 2A).

**Figure 2.**
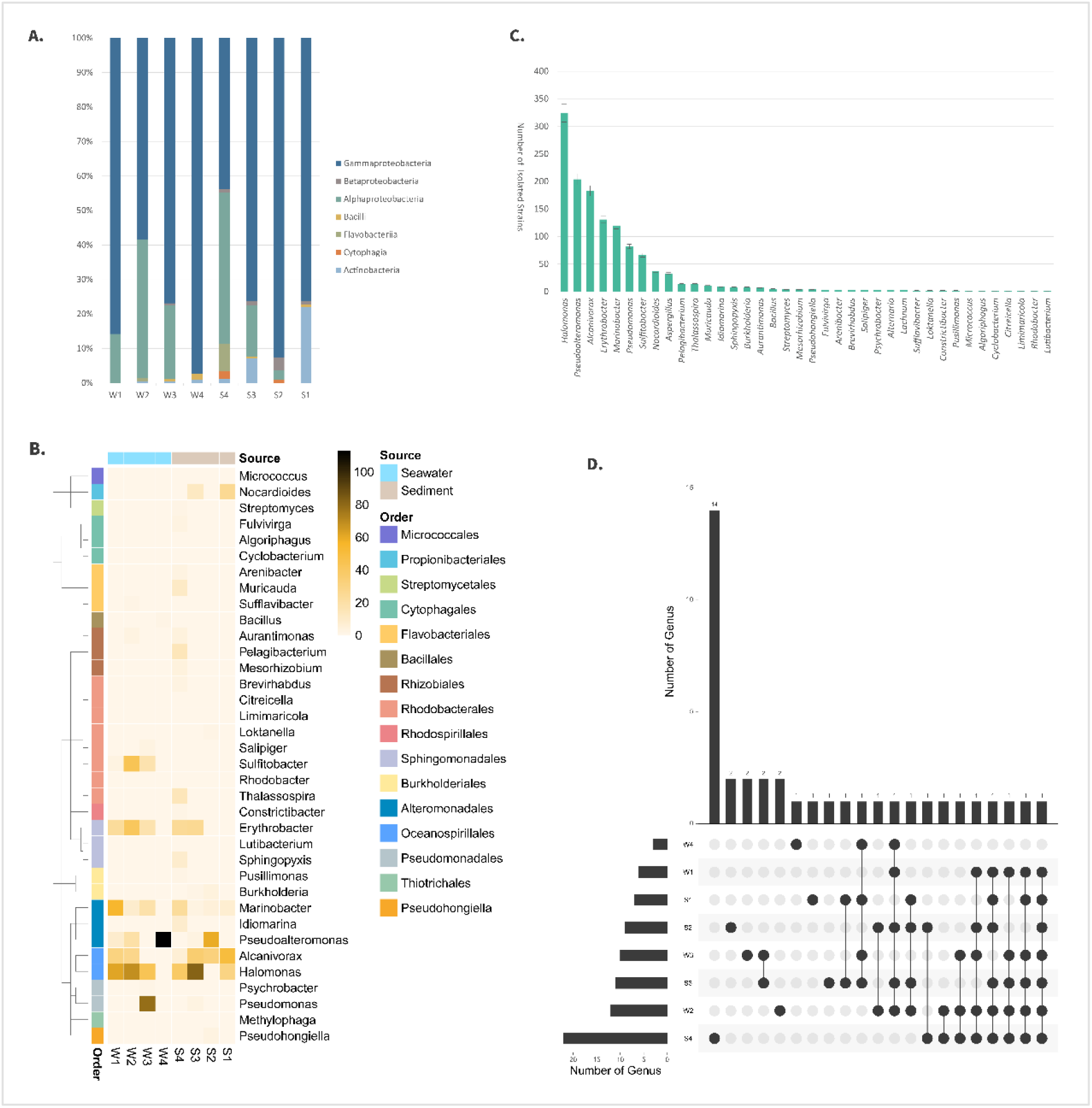
Microbial diversity of culturable bacteria in each sample. (A) Diversity of culturable bacteria in each sample at class level; (B) Diversity of culturable bacteria in each sample at genus level; (C) Number of culturable bacteria of each genus; (D) Number of genera isolated from different samples & number of genera unique and common to different samples.

At the genus level, *Halomonas* was distributed in all the samples, and its relative abundance was the highest in S3 and W2. *Alcanivorax* was also widely distributed, but the number of isolates was lower than *Halomonas*. For the seawater samples, the relative abundances of *Halomonas* and *Erythrobacter* showed a peak in W2 and a “low-high-low-extremely low” trend. The relative abundance of *Marinobacter* presented a “high-low-high-extremely low” trend. *Pseudoalteromonas* mostly concentrated in W4. For the sediment samples, the relative abundances of *Alcanivorax* and *Marinobacter* decreased with increasing depth; *Halomonas* showed no obvious trend, and its distribution of S3 site was high. *Erythrobacter* was mainly distributed in S3 and S4 sites. *Nocardioides* mainly distributed in S1 and S3 sites (Figure 2B). Most of the rare bacteria were distributed in S4 station. *Marinobacter* were distributed in 7 samples, *Halomonas* and *Alcanivorax* were distributed in 4 samples, *Erythrobacter* and *Sulfitobacter* were distributed in 5 samples. Combined with Figure 2C, it can be seen that the number of microorganisms isolated from the widely distributed genera is relatively high.

There were 22 bacterial genera from S4, which was collected from 11km below the seafloor, and there were 14 unique genera among them. There were only 4 genera isolated from W4, the 11km seawater sample, and there was only one unique genus named *Micrococus* in it (Figure 2D). As samples at the same depth, W4 and S4 showed significant differences in the number of isolated species, which may be due to the increasing pressure. The microbial diversity isolated from four sediment samples increased with depth, while the microbial diversity isolated from four seawater samples showed no obvious trend.

#### Diversity of culturable bacteria isolated from different media

The dominant bacteria phylum obtained from the 6 media were Proteobacteria, and the dominant class were γ-Proteobacteria. The dominant bacteria orders of 5 media were Oceanospirillales, except that the dominant bacteria of the MG+ medium was Alteromonadales. At the phylum level, 1/10 2216E+ medium showed better selectivity to Bacteroidetes, while at the class level it showed better selectivity to Flavobacteria. AIA+ medium showed better selectivity to Proteobacteria at the phylum level and β-Proteobacteria at the class level. MG-medium showed better selectivity to Actinobacteria at the phylum level and Actinobacteria at the class level (Figure 3A, B). **In brief, media type had an effect on microbial diversity of water and sediments at phylum and class level**. At genus level, the bacterial diversity of the 6 media from high to low was 1/3 R2A + (24 genera), MG-(21 genera), 1/10 2216E+ (21 genera), MG+ (18 genera), CDB+ (17 genera), and AIA+ (14 genera). There are some differences in the higher taxa, but the general trend is the same (Figure 3C).

**Figure 3.**
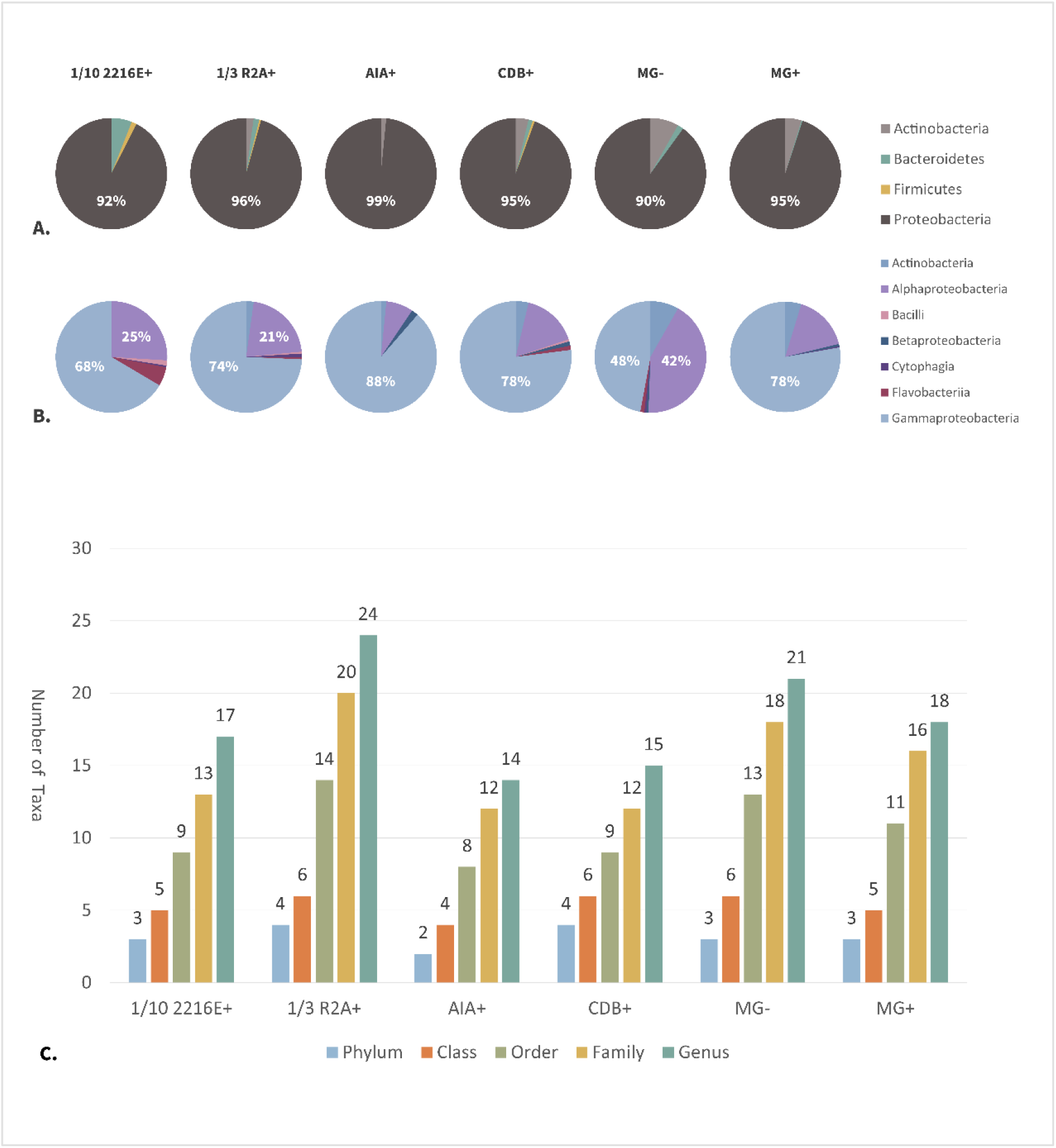
Microbial diversity of culturable bacteria in each medium. (A, B) Community structure of culturable bacteria isolated from different media at phylum and class levels; (C) Diversity of culturable bacteria isolated from different media at various taxonomic levels

The dominant bacteria isolated from each medium was *Halomonas*, showing its strong environmental adaptability. *Pseudoalteromonas* was the second dominant bacteria in AIA+ and MG+ medium. *Alcanivorax* was the second dominant bacteria in 1/3 R2A+ medium. In the medium of MG- and 1/10 2216E+, *Erythrobacter* was the second dominant genus. In CDB +, the second dominant genus was *Marinobacter*.

1/10 2216E+ showed strong selectivity to *Muricauda*, certain selectivity to *Bacillus*, and enrichment to *Erythrobacter*. 1/3 R2A+ showed selectivity to *Fulvivirga, Aurantimonas, Citreicella* and *Sphingopyxis*, and enrichment to *Alcanivorax*. AIA+ showed selectivity to *Burkholderia*, enrichment to *Pseudoalteromonas* and *Halomonas*, CDB+ showed selectivity to *Salipiger*. MG-showed selectivity to *Arenibacter, Mesorhizobium, Pseudohongiella* and *Nocardioides*, enrichment to *Sulfitobacter*, and MG+ showed selectivity to *Pelagibacterium*, and it has a strong enrichment effect on

#### *Marinobacter*. (Figure 4A, B)

9 genera could be isolated from all media. There were 24 genera isolated from 1/3 R2A+ and 3 unique genera isolated from 1/3 R2A+. There were only 14 genera isolated by AIA+ and no unique genera. Different media showed selectivity to different bacterial genera. Except for AIA+, the other five media could all cultivate microorganisms that could not be cultured in other media. There were 4 unique genera isolated from MG-, 3 unique genera isolated from 1/3 R2A+, 2 unique genera isolated from 1/10 2216E+, and 1 unique genus isolated from CDB+ and MG+ (Figure 4C). Unique genera of culturable bacteria isolated from different media were shown in Table 2.

**Table 2.** Unique genera of culturable bacteria in different media.

| **Medium type** | **The unique genus isolated from each medium** |
| --- | --- |
| 1/10 2216E+ | Lutibacterium, Cronobacter |
| 1/3 R2A+ | Citricella, Rhodobacter, Methylophaga |
| AIA+ | none |
| CDB+ | Salipiger |
| MG- | Micrococcus, Algoriphagus, Mesorhizobium, Limimarinicola |
| MG+ | Cyclobacterium |

**Figure 4.**
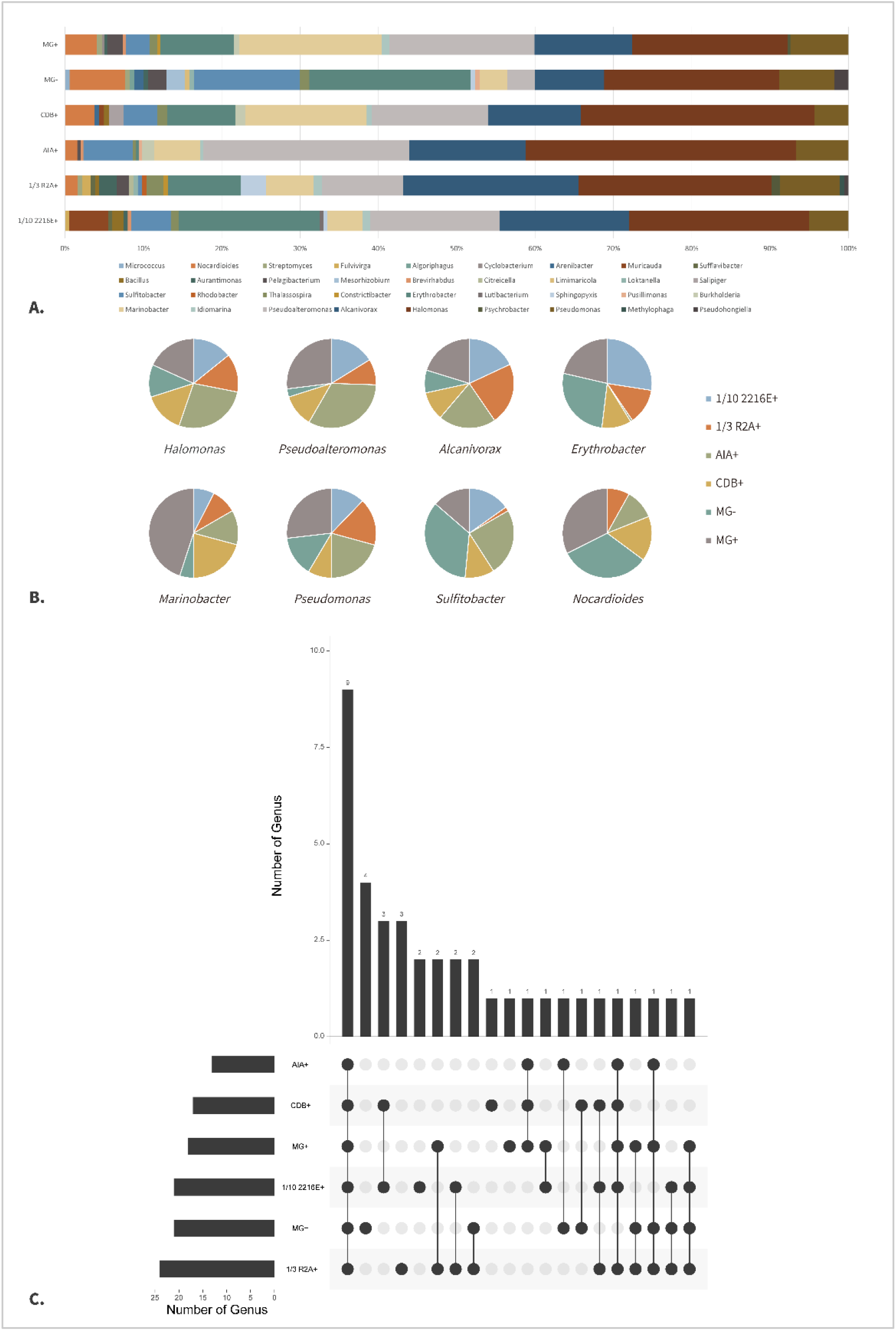
Relative abundance of culturable bacteria and selectivity of media. (A) Relative abundance of culturable bacteria isolated from different media at genus level; (B) Selectivity of the 6 media to the top 8 genus (C) Number of genera isolated from different media & number of genera unique and common to different media

#### Diversity of culturable bacteria isolated from the deepest sample using more kinds of media

Based on the diversity data obtained previously, 9 new media were added for further microbial isolation culture in the sample S4, which was from the deepest site and had the largest microbial diversity.

877 strains of microorganisms were obtained, belonging to 5 phyla, 11 classes, 20 orders, 23 families and 28 genera. Compared with the former result in S4, 1 new phylum, 6 classes, 10 orders, 10 families and 19 genera were isolated. Among them, 1 phylum, 6 classes, 8 orders, 8 families and 14 genera have not been isolated even in other sites (Figure 5).

**Figure 5.**
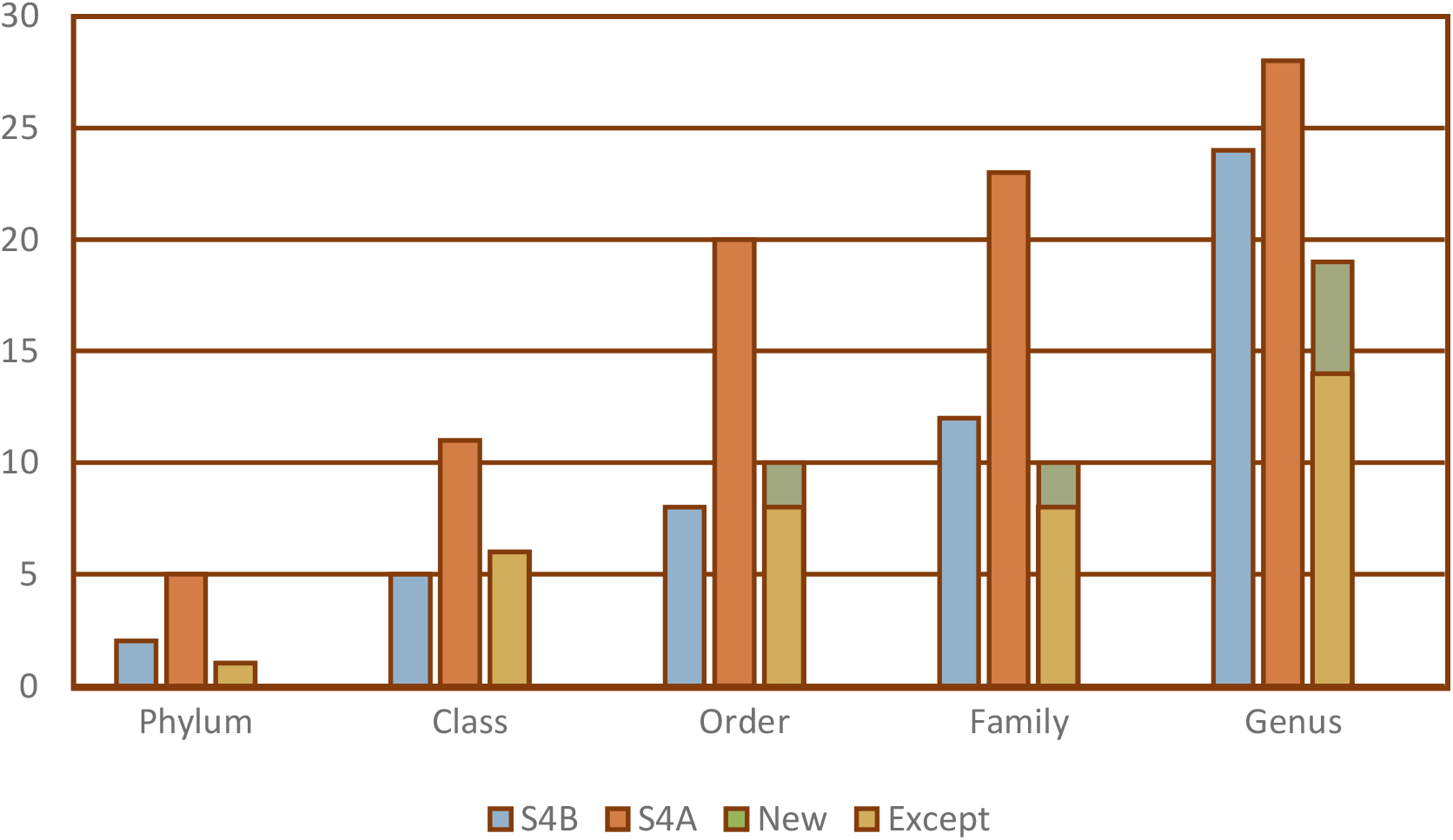
Microbial diversity at each classification level before and after adding extra culture media.

Several special media had certain selectivity for specific strains. Five strains of Flavobacteriaceae were obtained from SCA medium, accounting for 13.9% of the strains. 2 strains of Actinomycetes belonging to *Microbacterium* and *Streptomyces* were isolated from AIA. At the genus level, one strain of *Alcanivorax* and one strain of *Yangia* were isolated from nutrient medium R+. 3 strains of *Bacillus* were isolated from 1/10 2216E+ medium, AIA medium and R+ medium respectively. One *Sulfitobacter* strain was obtained from AIA and PDA+, respectively.

By comparison, it was found that adjusting salt concentration (30g/L) in common media could significantly change the number and diversity of microorganisms obtained. In this study, four common culture media R, PDA, AIA and G were modified with salt concentration adjustment. The results showed that the number of strains and the number of genus groups increased after R and AIA culture media were modified, showing higher diversity, and there were significant differences in the population structure of culturable microorganisms.

### Potential novel bacterial species from the 8 samples

25 bacterial strains (1.9% of the total) showed 16S rRNA gene similarities of less than 98.65% to the type strains of their closest known bacterial species, and represented 16 potential novel species. These strains were isolated from different depths, including ~ 1000 m (2 strains), ~ 4000 m (6 strains), ~ 6000 m (5 strains), ~ 6500 m (7 strains), ~ 7000 m (1 strain) ~11000 m (4 strains); no strains of potential novel species were obtained from W4. Moreover, 9 of the 25 potential novel strains were obtained from MG+ medium, 6 strains were obtained from MG-, 5 strains were obtained from AIA+, 4 strains were obtained from CDB+, 1 strain was obtained from 1/3 R2A+. These potential novel strains belonged to α-Proteobacteria (3 strains), β-Proteobacteria (3 strains), γ-Proteobacteria (17 strains), Actinobacteria (1 strain) and Bacilli (1 strain).

### The difference of environmental organic matter in each sample

Four sediment samples were tested for environmental organic matter using untargeted LC-MS. There were more than 11,000 cationic mass spectra peaks. A total of 340 substances were annotated, and 285 of them were in public databases such as HMDB and Lipidmaps, while 107 metabolites were annotated in KEGG database.

S1 was significantly different from the samples at the other three sites. The metabolites in S1 were separated from others at the subcluster level (Figure 6A, B). The content of Laccarin, 5’-Deoxyadenosine, L-Valine, and Dihydrocoumarin in S2, S3, S4 varied from each other, but there was no significant difference in other metabolites (Figure 6C).

**Figure 6.**
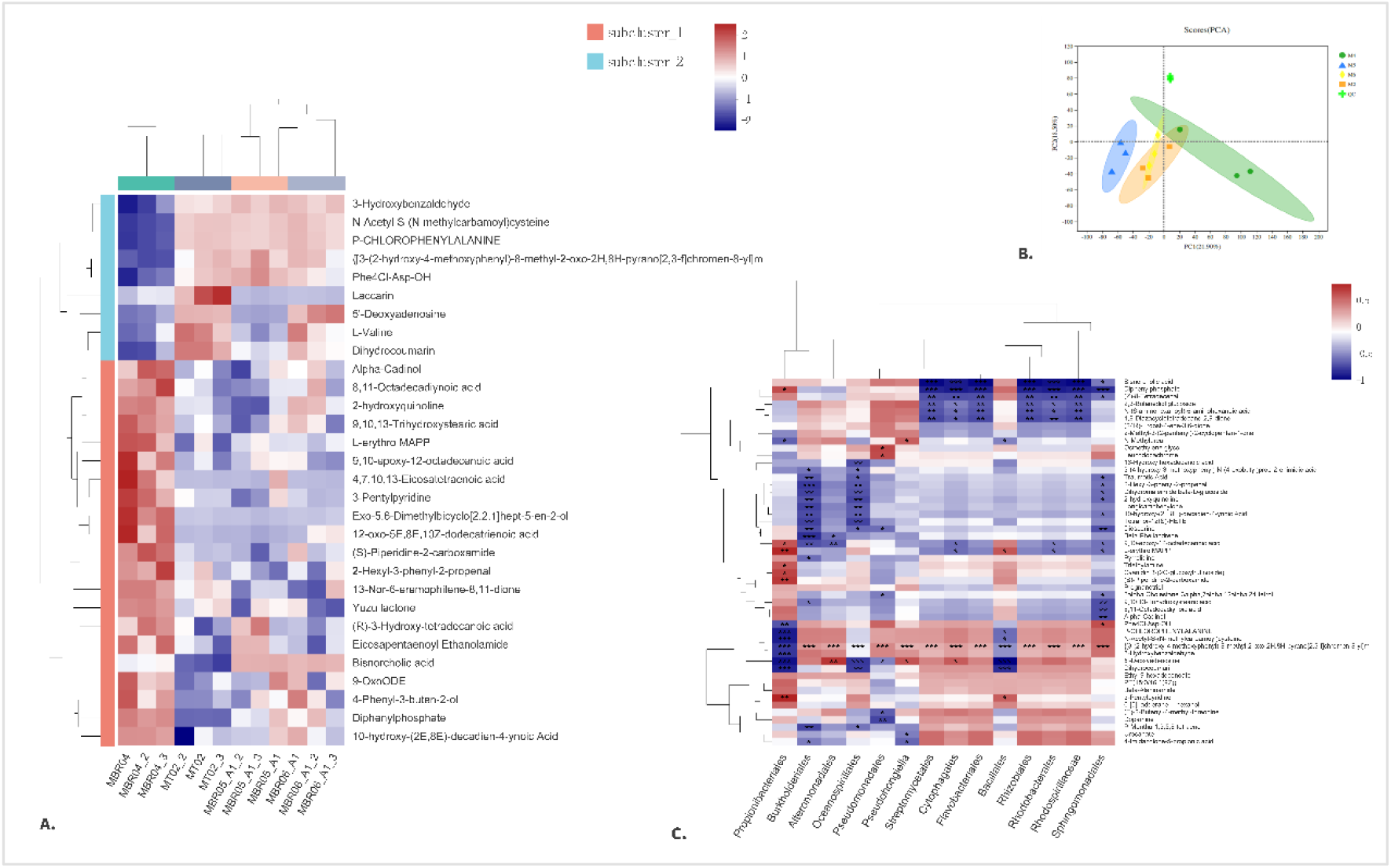
Analysis of environmental organic matter. (A) Hierarchical cluster heat map of environmental organic matter; (B) Principal component analysis of environmental organic matter of samples; (C) Correlation analysis between microbial diversity and environmental organic matter.

### Correlation analysis between microbial diversity and environmental factors

In order to verify the relationship among environmental parameters, microbial diversity and environmental organic matter diversity, Mantel test and Partial Mantel test were carried out for each matrix. The Mantel tests on the sample-environmental parameter matrix and the sample-microbial diversity matrix showed that the p-values were all less than 0.05 at phylum, class, order, family and genus levels, and there was a significant correlation between the the two matrices, while the R values of Mantel tests ranged from 0.2932 to 0.7716.. While the Partial Mantel test results of removing environmental organic matter showed that at the phylum, class, order and genus levels, there was a significant correlation between the two matrices. **In other words, environmental parameters can explain the diversity of culturable microorganisms at different stations**.

The Partial Mantel tests excluding the influence of environmental parameters for sample-microbial diversity matrix and sample-environmental organic matter diversity matrix showed that there was a significant correlation between the sample - microorganism diversity matrix and the sample - metabolite diversity matrix. **That is, at the phylum, class, and family levels, environmental organic matter shape the culturable bacterial diversity**.

## Discussion

### Culturable microbial diversity in the Mariana Trench

In this study, strains of bacteria isolated from the Mariana Trench were classified into Actinobacteria, Bacteroidetes, Firmicutes, Proteobacteria. The majority of microorganisnm were Proteobacteria and Firmicutes, while Actinobacteria and Bacteroidetes were in the minority, which was consistent with the recent high-throughput sequencing results of the sediments in the southern Mariana Trench. However, in this study, the relatively low abundance of Cyanobacteria, Planctomycetes, Chloroflexi, some Archaea and Methanogens were not isolated, which may be related to the special requirements of these microorganisms on the medium.

At the genus level, *Halomonas* and *Pseudoalteromonas* were the dominant bacteria, accounting for 25.59% and 16.11% respectively, which is different from the results of Huang Ying *et al* (). Huang’s study showed that *Sulfitobacter* was the main microorganism isolated from the seawater sample of S4, followed by *Pseudoalteromonas*. In our study, *Halomonas* was the main microorganism while *Sulfitobacter* accounted for a low proportion (5.21%).

The metagenome revealed the presence of Cyanobacteria in S4 (Chen P, Zhou H, Huang Y, et al. Revealing the full biosphere structure and versatile metabolic functions in the deepest ocean sediment of the Challenger Deep. Genome Biol. 2021,22(1):207.). Generally, Cyanobacteria has strong environmental adaptability and is easy to cultivate, but it wasn’t cultivated in our study. It can be refered that the existence of Cyanobacteria in the metagenome is due to the deposition of fragments of Cyanobacteria from the photic zone, which interferes with the metagenomic results and makes it impossible to accurately reconstruct the true survival state of microorganisms in the deep sea sediments, so it is necessary to isolate the culturable microorganisms.

The five widely distributed taxa we isolated are *Marinobacter, Halomonas, Alcanivorax, Erythrobacter, Sulfitobacter. Marinobacter* is a kind of halophilic and diazotrophic bacteria, which is capable of decomposing hydrocarbons. It’s especially suitable for bioremediation of hydrocarbon pollutants in a high-salt environment without enough nitrogen compounds. The whole genome sequencing of one strain, *Marinobacter Flavimaris* LMG 23834T, also revealed its potential for biological repair. In addition, the existence of haloduric or halophilic bacteria like *Halomonas* provides a theoretical possibility for biological treatment of wastewater with high salinity. Some studies have also shown that denitrification capacity is an important phylogenetic molecular marker for distinguishing different species of *Halomonas. Alcanivorax* is often used for biodegradation of large molecules, like polycyclic aromatic hydrocarbons. *Erythrobacter* is widely distributed and has the ability to decompose aromatic compounds, while some individuals also have the ability to produce carotenoids and cytotoxins. Some *Sulfitobacter* bacteria are thought to be related to microalgal metabolism, such as secreting IAA to stimulate diatoms to divide, and the synthesis of IAA depends on endogenous tryptophan produced by diatoms. In a word, the strains isolated in the study has great potential for research, development and application.

### Medium effect and improvement

1/10 2216E+ and 1/3 R2A+ were diluted and adjusted according to the chemical composition of the sediment in the sampling area and previous research experience. While AIA+, MG, MG+ and CDB+ were adjusted based on the preliminary experimental results. It is worth noting that both *Halomonas* and *Alcanivorax* belong to the order of Oceanospirillales, and are close relatives, along with microorganisms of the genera *Pseudoalteromonas, Erythrobacter*, and *Marinobacter*, these groups have been reported to have the ability to degrade refractory organic compounds or have the potential metabolic pathway of microbial syntrophy. Such ability to use nutrients efficiently may be the reason why they tend to become the dominant genera.

As the most commonly used medium for the isolation of seawater microorganisms, 2216E has been widely used in laboratory. R2A itself is a medium with low nutrient concentration,which is often used to isolate microbes from environmental samples. In this experiment, 1/10 2216E+ and 1/3 R2A+ were used as nutrient adjustment medium, which indeed achieved better separation effect than other medium with higher nutrient concentration. We found that 1/10 2216E+ medium had the highest abundance of microorganisms, which was obviously not at the same order of magnitude as other media, but the separation efficiency of 1/10 2216E+ medium was not as good as that of 1/3 R2A+ medium at genus level. Literature and research experience suggest that 1/10 2216E+ medium has certain universality, but it is not the best choice for experiments aiming at higher diversity of isolation. This also provides some ideas for the isolation of microorganisms in similar environments.

All media in this study showed some selectivity for some rare groups of microorganisms. We take 1/10 2216E+ medium as an example, which has a strong selection effect on the bacteria of genus *Muricauda*. There are few studies on *Muricauda*, most of which focus on its co-culture with microalgae and Cyanobacteria. Studies have shown that when *Muricauda* and microalgae are co-cultured under mixed nutrition, they can significantly promote the growth of microalgae, and the increase of microalgal cell density is often accompanied by the increase of *Muricauda* cell number. One strain of *Muricauda* has been shown to contain appendages with fibrous vesicles on the surface. *Muricauda* to connect cells to each other or interact with the surrounding environment. A study on the response of autotrophic and heterotrophic microorganisms to temperature showed that temperatures higher than 34°C were unfavorable for Cyanobacteria and *Muricauda* in co-culture. There are more peptone, carbonate, strontium, citrate in 1/10 2216E+ medium than other media, and perhaps these substances is what *Muricauda* needs to grow. Given that there are Cyanobacteria in the metagenome results, if the presence of *Muricauda* can promote the growth of Cyanobacteria, and we isolated *Muricauda* but did not isolate Cyanobacteria, maybe we can further speculate that there is no active Cyanobacteria in the deep sea.

In short, the isolation of these special strains is the prerequisite for future research and application. If the culture media can be further improved on the basis of our research to enrich the microorganisms of these rare groups, it will promote the culture of microorganisms of rare groups. It can also be further demonstrated that the microorganisms adapted to the deepsea oligotrophic environment are indeed more sensitive to nutritional conditions.

### Impact factor

We found that environmental nutrients do have a significant effect on culturable microbial diversity. A previous study has shown that the possible provenances of the sinking particles of the Mariana Trench Challenger Deep include volcanic sources, eolian dust terrestrial and marine authigenic minerals and the volcanic sources are the dominant provenance and consist mainly of the surrounding ridges and sea basins, among which the West Mariana Ridge and the Kyushu-Palau Ridge contribute the most. The terrestrial materials and authigenic minerals brought by eolian dust contribute little. Moreover, the study shows there is an oxidative sedimentary environment in the Mariana Trench and that the oxidation degree of seawater increases with the depth. Therefore, this general oxidation environment may lead to the metabolic particularity of deepsea microorganisms, and the increase of oxidation degree with water depth may also result in the difference of microbial distribution at different depths.

In terms of inorganic ions, the samples contained N, P, S and a variety of trace elements. According to the results of environmental organic matter, there are many kinds of nutrients in the samples, and even many of them are unknown to us, so it can be considered to add or subtract relevant substances in the culture media to assist the separation and culture of microorganisms. Combined with the results of metagenomics, the microbes in the samples may have certain chemical autotrophic ability, fermentation ability, or can participate in N and S cycles.

In a word, differences in the diversity of culturable microorganisms isolated by us can be explained to some extent by differences in station position, and these microorganisms have certain metabolic potential. The isolation method, the direction of medium improvement and the correlation between the obtained environmental organic matter and diversity can further guide the optimization of the isolation method/culture conditions of deep-sea microorganisms.

## Supplemental material

**Table S1.** Media modified from several base-type that favored the growth of microorganisms.

| Media | Components/L |
| --- | --- |
| 1/10 2216E+ | Tryptone 0.5 g, Yeast extract 0.1 g, Ferric citrate 0.01 g, MgCl<sub>2</sub> 0.598 g, Na<sub>2</sub>SO<sub>4</sub> 0.324 g, CaCl<sub>2</sub> 0.18 g, KCl 0.055 g, Na<sub>2</sub>CO<sub>3</sub> 0.016 g, KBr 0.008 g, SrCl<sub>2</sub> 0.0034 g, H<sub>2</sub>BO<sub>3</sub> 0.0022 g, Na<sub>2</sub>SiO<sub>4</sub> 0.0004 g, NaF 0.00024 g, NH<sub>4</sub>NO<sub>3</sub> 0.00016 g, Na<sub>2</sub>HPO<sub>4</sub> 0.0008 g, KNO<sub>3</sub> 0.004 g, NaCl 31.945 g, Agar 10g (to prepare a solid medium) |
| 1/3 R2A+ | Yeast extract 0.17 g, Casein Hydrates 0.17 g, Glucose 0.17 g, Soluble starch 0.17 g, K<sub>2</sub>HPO<sub>4</sub> 0.1 g, MgSO<sub>4</sub> 0.008 g, Sodium pyruvate 0.1 g, KNO<sub>3</sub> 0.004 g, NaCl 30 g, Agar 10g (to prepare a solid medium) |
| AIA+ | Sodium caseinate 2 g, L-aspartic acid 0.1 g, CH<sub>3</sub>CH<sub>2</sub>COONa 4 g, K<sub>2</sub>SO<sub>4</sub> 0.5 g, MgSO<sub>4</sub> 0.1 g, FeSO<sub>4</sub> 0.001 g, NaCl 30g, Agar 10g, pH 8.1±0.2 |
| MG | KNO<sub>3</sub> 1.0g, KH<sub>2</sub>PO<sub>4</sub> 0.5g, MgSO<sub>4</sub> 0.5g, FeSO<sub>4</sub> 0.01g, NaCl 0.5g, Soluble starch 20.0g, Agar 10g (to prepare a solid medium), pH 7.2 - 7.4. **After sterilization and cooling to 45-55°C, add 1ml of 3% KMnO<sub>4</sub> solution to every 300mL medium.** |
| MG+ | KNO<sub>3</sub> 1.0g, KH<sub>2</sub>PO<sub>4</sub> 0.5g, MgSO<sub>4</sub> 0.5g, FeSO<sub>4</sub> 0.01g, NaCl 30.5g, Soluble starch 20.0g, Agar 10g (to prepare a solid medium), pH 7.2 - 7.4. **After sterilization and cooling to 45-55°C, add 1ml of 3% KMnO<sub>4</sub> solution to every 300mL medium.** |
| CDB+ | NaNO<sub>3</sub> 3.0g, K<sub>2</sub>HPO<sub>4</sub> 1.0g, MgSO<sub>4</sub> 0.5g, KCl 0.5g, FeSO<sub>4</sub> 0.01g, Sucrose 30.0g, NaCl 30.0g, Agar 10g (to prepare a solid medium), pH 5.8 - 6.2 |

**Table S2.**
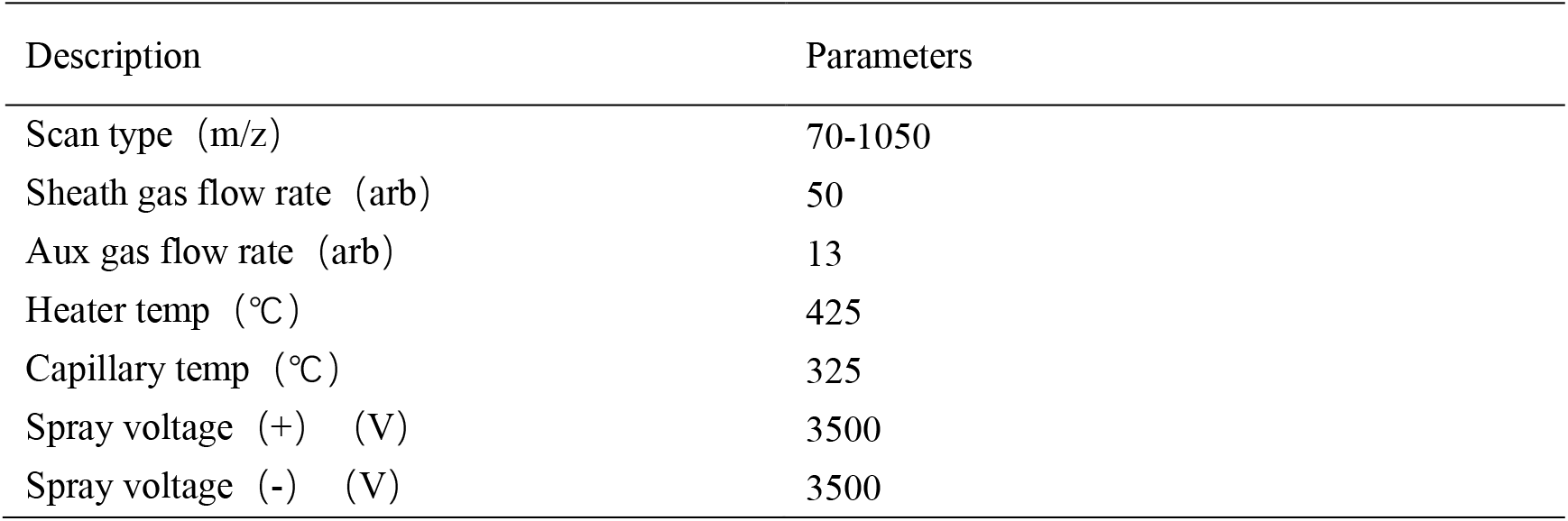
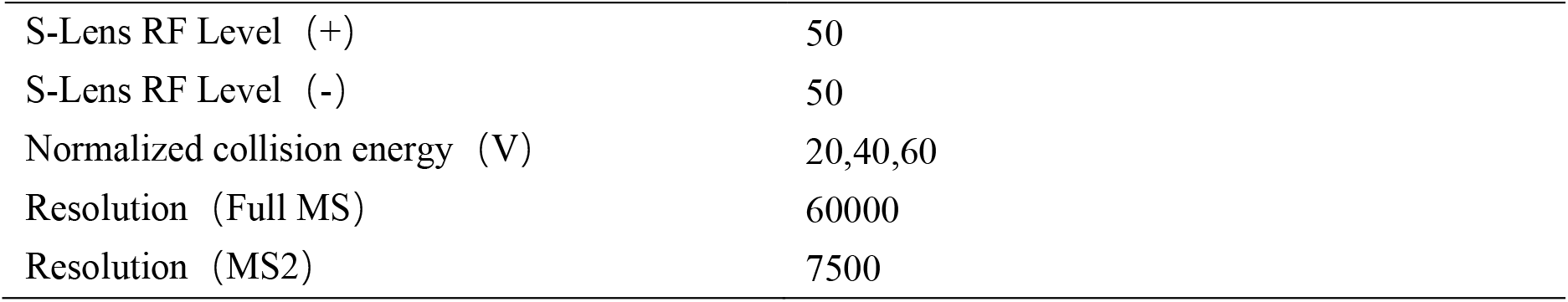
Multiple reaction monitoring (MRM) acquisition parameters.

| Description | Parameters |
| --- | --- |
| Scan type (m/z) | 70-1050 |
| Sheath gas flow rate (arb) | 50 |
| Aux gas flow rate (arb) | 13 |
| Heater temp (°C) | 425 |
| Capillary temp (°C) | 325 |
| Spray voltage (+) (V) | 3500 |
| Spray voltage (-) (V) | 3500 |
| S-Lens RF Level (+) | 50 |
| S-Lens RF Level (-) | 50 |
| Normalized collision energy (V) | 20,40,60 |
| Resolution (Full MS) | 60000 |
| Resolution (MS2) | 7500 |

